# Biological Correlations and Confounding Variables for Quantification of Retinal Ganglion Cells Based on Optical Coherence Tomography using Diversity Outbred Mice

**DOI:** 10.1101/2020.12.23.423848

**Authors:** Adam Hedberg-Buenz, Kacie J. Meyer, Carly J. van der Heide, Wenxiang Deng, Kyungmoo Lee, Dana A. Soukup, Monica Kettelson, Danielle Pellack, Hannah Mercer, Kai Wang, Mona K. Garvin, Michael D. Abramoff, Michael G. Anderson

**Author notes:** Co-Corresponding: Dr. Michael G. Anderson, Department of Molecular Physiology and Biophysics, 3123 Medical Education and Research Facility, 375 Newton Road, Iowa City, IA 52242. Disclosure: **A. Hedberg-Buenz**, none; **C.J. van der Heide**, none; **K.J. Meyer**, none; **D. Soukup**, none; **W. Deng**, none; **K. Lee**, none; **H. Mercer**, none; **M. Kettelson**, Digital Diagnostics, Inc.; **D. Pellack**, none; **K. Wang**, none; **M.K. Garvin**, is the co-inventor on a US patent related to the approach used to segment retinal layers; she has personally waived all financial rights to said patent, but the University still has rights; **M.D. Abramoff**, is the inventor on patents and patent applications of artificial intelligence and machine learning algorithms for diagnosis and treatment and is a Founder, CEO, employee of, and investor in Digital Diagnostics Inc, Coralville, Iowa, USA.; and **M.G. Anderson**, none.

## Abstract

**Purpose:** Despite popularity of optical coherence tomography (OCT) in glaucoma studies, it’s unclear how well OCT-derived metrics compare to traditional measures of retinal ganglion cell (RGC) abundance. Here, Diversity Outbred (J:DO) mice are used to directly compare ganglion cell complex (GCC)-thickness measured by OCT to metrics of retinal anatomy measured ex vivo with retinal wholemounts or optic nerve cross sections.

**Methods:** J:DO mice (*n* = 48) underwent OCT and fundoscopic exams, with GCC-thickness measured using automated segmentation. Following euthanasia, RGC axons were quantified from para-phenylenediamine-stained optic nerve cross sections and RGC somas from BRN3A-immunolabeled retinal wholemounts with total cellularity assessed by TO-PRO or hematoxylin nuclear staining.

**Results:** J:DO tissues lacked overt disease. GCC-thickness (62.4 ± 3.7 µm), RGC abundance (3,097 ± 515 BRN3A^+^ nuclei/mm^2^; 45,533 ± 9,077 axons), and total inner retinal cell abundance (6,952 ± 810 nuclei/mm^2^) varied broadly. GCC-thickness correlated significantly to RGC somal density (r = 0.46) and axon number (r = 0.49), whereas total inner retinal cellularity did not. Retinal area (20.3 ± 2.4 mm^2^) and optic nerve (0.09 ± 0.02 mm^2^) cross-sectional area varied widely. Sex did not significantly influence any of these metrics. In bilateral comparisons, GCC-thickness (r = 0.89), inner retinal cellularity (r = 0.47), and RGC axon abundance (r = 0.72) all correlated significantly.

**Conclusions:** Amongst outbred mice with widely variable phenotypes, OCT-derived measurements of GCC thickness correlate significantly to RGC abundance and axon number. The extensive phenotypic variability exhibited by J:DO mice make them a powerful resource for studies of retinal anatomy using quantitative genetics.

## Introduction

Retinal ganglion cells (RGCs) are the first neuron in the visual phototransduction circuit to fire an action potential; input from RGC dendrites is summated at the soma and projected toward the lateral geniculate nucleus in the brain via RGC axons where it is ultimately processed into vision. Thus, RGCs are critical to the physiology of vision and damage to any part of these neurons is potentially threatening to sight. RGCs are post-mitotic, so their loss is irreversible and leads to permanent vision loss. The most common ophthalmic disease of RGCs is glaucoma^1^, but changes indicative of RGC damage are also a feature of several diseases with broader systemic effects, including diabetes^2^, Alzheimer’s disease^3^, multiple sclerosis^4, 5^, and some forms of traumatic brain injury^6, 7^, among others^8^. Clinically assessed structural indices of RGC damage in glaucoma have traditionally focused on features of optic nerve head morphology, such as increasing cup-to-disc ratio and the presence of notching. More recently, advances in optical coherence tomography (OCT) have made it practical to more broadly assess additional features of RGC loss, including thickness of the neuroretinal rim and thickness of the ganglion cell complex (GCC; combined thickness of areas containing the RGC axons, soma, and dendrites)^9–11^.

Although OCT-derived measurements are increasingly popular metrics for RGC disease—in both humans^12, 13^ and animal models^14^—OCT is prone to known artifact and some interpretations remain unclear^15, 16^. One key question that has not been fully addressed is whether non-invasive OCT-derived measurements are in fact associated with biological measurements of RGC abundance made from histology-based analyses, such as quantifications from retinal wholemounts or cross sections of the optic nerve. Here, we contribute to the understanding of this issue by using mice to study the relationships of in vivo OCT-derived ganglion cell complex (GCC) thickness to ex vivo quantified metrics of RGC abundance. RGC number is influenced by heredity as a complex trait^17–19^, so we utilized outbred J:DO mice for these experiments. J:DO mice are a stock of mice with a high degree of genetic heterogeneity, and therefore expected to have a corresponding wide range of RGC numbers between individuals. Our overall experimental design involved acquiring a large cohort of J:DO mice, non-invasively imaging all retinas using OCT, and using automated segmentation to quantify GCC thickness. Following euthanasia, we subsequently performed histology and/or immunostaining coupled with semi-automated quantifications of total cellular and RGC density in the inner retina and RGC axon number in the optic nerve. From these data, we were able to evaluate biological correlates of GCC thickness, as well as gain insight into previously unknown aspects of basic ocular anatomy and potential confounding variables relevant to RGC quantification.

## Methods

### Experimental Animals

Mice were maintained on a 4% fat NIH 31 diet provided *ad libitum*, housed in cages containing dry bedding (Cellu-dri; Shepherd Specialty Papers, Kalamazoo, MI), and kept in a 21°C environment with a 12-h light: 12-h dark cycle. All mice were treated in accordance with the Association for Research in Vision and Ophthalmology Statement for the Use of Animals in Ophthalmic and Vision Research. All experimental protocols were approved by the Institutional Animal Care and Use Committee of The University of Iowa.

### Sample Numbers

Data were collected from all mice in an identical order, but implementation of inclusion/exclusion criteria and attrition resulted in losses such that the number of samples utilized in final data sets were not equal across all assays (Summarized in Supplemental Fig. 1). A cohort of adult Diversity Outbred mice^20^ (J:DO; *n*=48 mice, equal numbers of males and females, 8 weeks of age) were ordered from The Jackson Laboratory (Stock no: 009376, Bar Harbor, ME) and subsequently housed at the University of Iowa Research Animal Facility. A total of 15 mice died between 8 and 20 weeks of age, frequently related to anesthesia but also without apparent cause during aging; tissues from these mice were excluded from all immunohistochemical and histologic analyses. At 16 weeks of age, both eyes of 47 mice underwent OCT imaging. Scans from two mice failed quality control when later subjected to automated segmentation (described in detail below); data from both eyes of these mice were excluded from the GCC thickness dataset (leaving 45 pairs of eyes in the final GCC thickness dataset). Tissues from the two mice excluded from the GCC dataset were none-the-less carried through and contributed to the immunohistochemical and histologic analyses. One to two days following OCT imaging, the right fundus of 47 mice was imaged by fundoscopy, with one mouse (#15725) having developed a cloudy cornea after OCT imaging that prevented imaging (leaving 46 images in the final fundoscopy data set); mouse #15725 with the cloudy cornea was later found to also have a damaged optic nerve, thus the retina and optic nerve from this apparently induced unilateral anomaly were excluded. At 20 weeks of age, there were 33 mice which were euthanized, and tissues collected. From 33 retinas available for the left eye analysis, four were damaged in the original flat-mounting procedure and one (from mouse #15725) was excluded as explained above (leaving 28 retinas in the final BRN3A dataset). For unknown reasons, one of the BRN3A-labeled retinas was insufficiently stained by TO-PRO (leaving 27 retinas in the final TO-PRO dataset). When these same retinas were later stained with H&E another six succumbed to stresses of processing (i.e. retinas became delaminated and/or fractured into smaller pieces) and were unable to be analyzed (leaving 22 retinas in the left eye H&E dataset). From 33 retinas available for the right eye analysis, six became damaged at the time of flat-mounting (leaving 27 in the final right eye H&E dataset). From 66 optic nerves, both nerves of 2 mice were inadvertently lost during dissection or processing, and 1 (from mouse #15725) was excluded as explained above (leaving 61 optic nerves, 30 pairs, in the final PPD dataset). When data was excluded for the reasons described above, it was done so by an investigator masked to other data for that sample; no samples were excluded based on the appearance of being an outlier in the final graphs.

**Figure 1.**
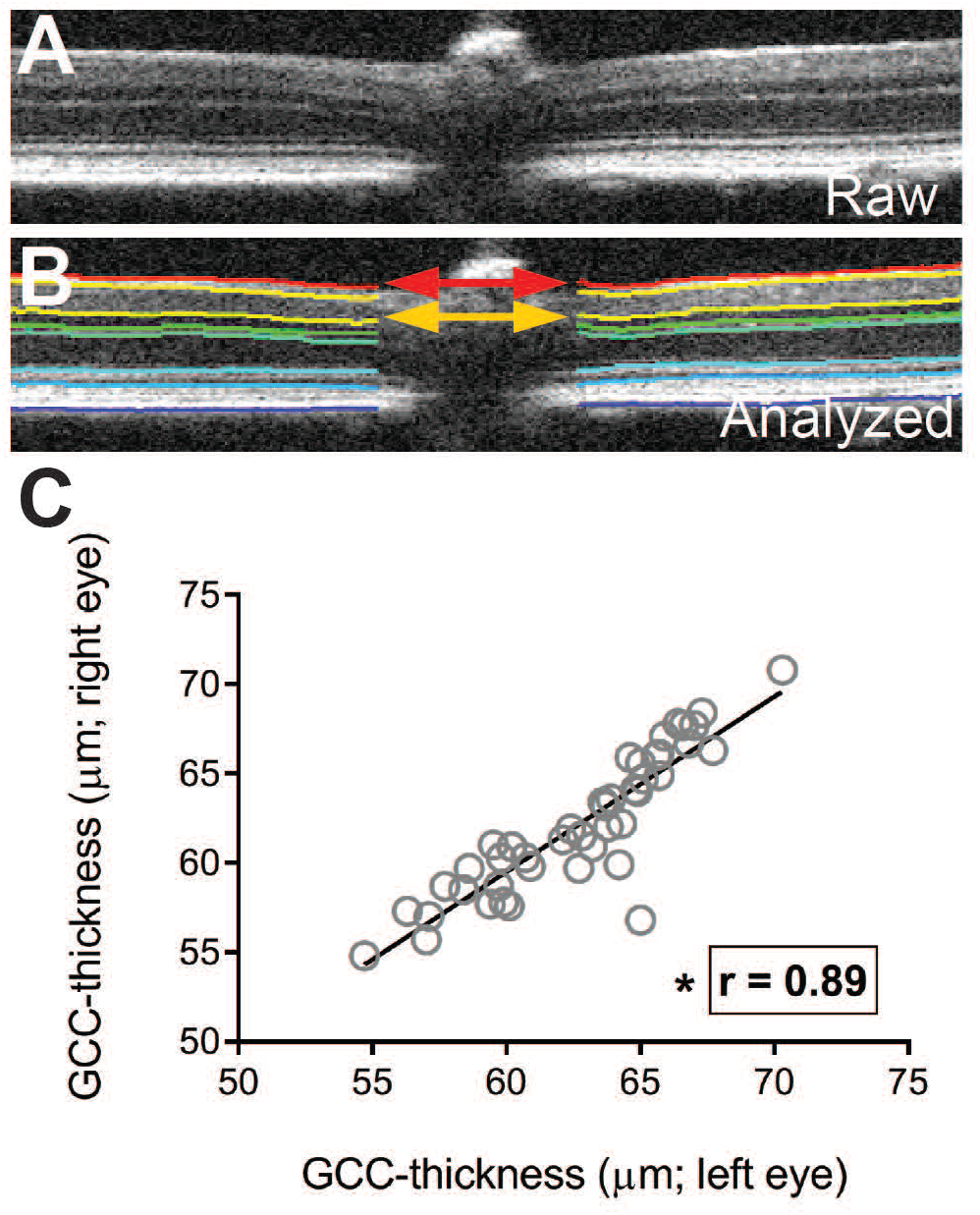
Retinal ganglion cell complex thickness is variable across individual J:DO mice but conserved within individuals. Image scans from in-vivo retinal scans obtained by optical coherence tomography (OCT), were analyzed using custom algorithms to segment and quantify thickness of retinal layers, and specifically the retinal ganglion cell complex (GCC). Correlation testing of GCC thickness between the left and right eyes of the same mouse within the study cohort of J:DO mice. A representative OCT-image of retina in (**A**) raw and (**B**) analyzed form. Following analysis of the raw image, the analyzed form contains inset lines to denote segmentation of the different layers (by differing color) within the retina. The GCC consists of the: nerve fiber, ganglion cell, and inner plexiform layers combined, and its thickness is the sum of these layers (vertical distance between red and gold arrows. (**C**) Graph plotting thickness of the right versus left GCC, showing a strong correlation (r = 0.89) of GCC thickness between eyes of the same individual in a cohort of J:DO mice. Each dot represents mean GCC-thickness of the right (*y-axis*) versus left (*x-axis*) eye from an individual J:DO mouse (*n*=45 mice, 90 retinas), inset line represents the best-fit line, Pearson’s correlation coefficient (r), and the asterisk represents a P *<* 0.05 using a two-tailed Student’s *t*-Test.

### Spectral domain optical coherence tomography (OCT) imaging and analysis

Mice were anesthetized with a mixture of ketamine (87.5 mg/kg; VetaKet, Akorn) and xylazine (12.5 mg/kg; AnaSed® Injection, Akorn) by intraperitoneal injection and corneas were kept lubricated with Balanced Salt Solution (BSS®; Alcon Laboratories, Fort Worth, TX). Anesthetized mice were placed onto an adjustable cassette connected to a platform to allow three-dimensional movement (referred to as “OCT” hereafter; Bioptigen Envisu R2200, Morrisville, NC). OCT scanning was centered on the optic nerve head of the retina and aligned in the horizontal and vertical planes^2^. OCT volume dimensions were 400 x 400 x 1024 voxels (1.4 x 1.4 x 1.566mm^3^). Following imaging, mice were administered yohimbine (2 mg / kg of body weight; Yobine® Injection, Akorn), provided supplemental indirect warmth for anesthesia recovery, and eyes were hydrated with ointment (Artificial Tears, Akorn), as described previously^21^.

Offline automated segmentation was performed using the Iowa Reference Algorithms 4.0 for three-dimensional automated layer segmentation; the thickness of the GCC was quantified as the distance between the upper surface of the inner limiting membrane (ILM) and bottom surface of the inner plexiform layer^22, 23^. An independent validation has shown favorable performance of this algorithm in comparison to other known algorithms for the inner retina in mice^24^. Quality control was performed on all segmentations using a series of objective exclusion criteria which led to the exclusion of scans from two mice. Image scans were omitted from analyses if three of the four objective metrics were satisfied: quality index (QI) < 11.9^25^, maximum tissue contrast index (mTCI) < 10.4^26^, (RetMinCost, unit less) > 33,181, duration of analysis (RetTime, in seconds) > 312, or presence of visually erroneous segmentations. Exclusion criteria cut-offs for these metrics were set at two standard deviations above (retMinCost, RetTime) or below (mTCI) the mean for all retinal scans.

### Fundus Imaging

Pupils were dilated using a combination of 2% cyclopentolate hydrochloride ophthalmic solution (Cyclogyl®, Alcon Laboratories, Fort Worth, TX) and 2.5% phenylephrine hydrochloride ophthalmic solution (Paragon BioTeck, Inc., Portland, OR). Once pupils were fully dilated, mice were anesthetized with a mixture of ketamine (87.5 mg/kg; VetaKet, Akorn) and xylazine (12.5 mg/kg; AnaSed® Injection, Akorn) by intraperitoneal injection and corneas kept lubricated with BSS® (Alcon Laboratories, Fort Worth, TX). Following anesthesia administration, hypromellose 2.5% ophthalmic demulcent solution (Goniovisc®, HUB Pharmaceuticals, LLC, Rancho Cucamnoga, CA) was applied to each eye. Right eyes were imaged with a Micron III retinal imaging microscope (Phoenix Research Labs, Pleasanton, CA). Following imaging, mice were administered yohimbine (2 mg / kg of body weight; Yobine®, Akorn), were provided supplemental indirect warmth for anesthesia recovery, and their eyes were hydrated with ointment (Artificial Tears, Akorn), as described previously^21^.

### Collection of retinas and optic nerves

Mice were euthanized by carbon dioxide inhalation with death confirmed by cervical spine dislocation. Retinas were collected as previously described^27^. In brief, eyes were collected and the posterior eye cups were dissected and drop fixed in ice-cold 4% paraformaldehyde in 1x phosphate buffered saline (PBS) for four hours, rinsed in PBS at 4°C, from which point, the left and right eyes from each mouse were prepared for hematoxylin-eosin staining or BRN3A immunolabeling methodologies, respectively. Optic nerves were collected and processed for histology as previously described^28^. In brief, heads were removed from mice and submerged into half-strength Karnovsky’s fixative for 24 hours and rinsed in 0.1 M sodium cacodylate at 4°C.

### Preparation, imaging, and quantitative analysis of hematoxylin-eosin stained retinas

Fixed retinas from the left eye of mice were processed, whole-mounted, stained with H&E, imaged, and quantitatively analyzed as previously described^29^. In brief, dissected and processed retinas were transferred to positively charged glass microscope slides, mounted flat, dried overnight, and stained with H&E. Two images in each concentric zone (peripheral, mid-peripheral, and central) for each of the four petals of each retina were collected (*n*=24 images per retina), using a light microscope (BX52; Olympus, Tokyo, Japan) with identical camera (DP72, Olympus, Tokyo, Japan) and software (CellSens; Olympus, Tokyo, Japan) settings. Before further processing, retinal image sets were screened for quality control. For inclusion in the study: 1) eight images had to be collected from each of the three zones of concentricity (peripheral, mid-peripheral, and central) of the retina, for a total of 24 images, and 2) after the removal of artifacts (including tears, holes, and debris) from images using the RetFM-J image analysis plugin^27^, the sum of analyzed area (termed *included area* in RetFM-J) from all 24 images combined had to exceed 85% (or 2.86 of the 3.36 mm^2^) of the total retinal area sampled. Samples damaged during processing, such that 24 non-overlapping and concentrically distributed sample areas could not be imaged, were excluded. Finally, the RetFM-J plugin was run using Fiji image analysis software^30^ to segment and quantify nuclei from retinal image sets. Nuclei counted by RetFM-J were calculated as mathematical averages of nuclei across all images per retina and were expressed in terms of density.

### Preparation, imaging, and quantitative analysis of BRN3A immunolabeled retinas

Fixed posterior eye cups from the right eye of mice were permeabilized with 0.3% Triton-X 100 in PBS (PBST) overnight at 37°C. Retinas were dissected from cups and further permeabilized at −80°C for 15 min. and thawed at room temperature for 30 min. All following steps were carried out at room temperature, unless otherwise noted. Retinas were dissected and blocked with 2% normal donkey serum in PBST for three hours or overnight at 4°C. Retinas were incubated with an anti-BRN3A antibody (C-20, 1:200; Santa Cruz Biotechnology, Dallas, TX) in PBS with 2% normal donkey serum, 1% Triton-X 100, and 1% DMSO. Retinas were rinsed in PBST, incubated with a donkey anti-goat Alexa488-conjugated secondary antibody (A11055, 1:200; Life Technologies, Madison, WI) in PBS with 5% normal donkey serum, 1% Triton-X 100, and 1% DMSO, rinsed in PBST with TO-PRO-3® (abbreviated TO-PRO; ThermoFisher Scientific), mounted with Aqua-Mount (Lerner, Pittsburgh, PA), and topped with a weighted coverslip to promote flat mounting.

Immunolabeled retinas from J:DO mice were imaged using confocal microscopy (LSM710, Zeiss, Germany). For each retina, images (1024×1024 px, 425.1 mm^2^ image area) were collected at 400X total magnification from non-overlapping fields at each zone of eccentricity of the inner retina (such that *n* = 12 images total; 4 central, 4 mid-peripheral, 4 peripheral). Retinal image sets (*n* = 12 images) were opened in Fiji image analysis software ^30^ and prepared for quantitative analysis by a stepwise method using tools available within Fiji. Each image set underwent background subtraction with rolling ball radius set to 35 pixels, smoothened, conversion to binary using Huang thresholding, object erosion with subsequent dilation, watershed thresholding, and the fill holes function. Finally, BRN3A^+^ nuclei were segmented using the analyze particles function with a size inclusion limit of (20-150 µm^2^) and circularity (0-1). Density data were presented as the mean density of BRN3A^+^ nuclei per mm^2^ ± SD for each retina. Counts of total BRN3A^+^ RGC number were calculated by multiplying average density with area measured within each zone of eccentricity and summing the products for all three zones for each retina, as previously described ^31^. Samples damaged during processing, such that 12 non-overlapping and concentrically distributed sample areas could not be imaged, were excluded.

### Preparation, imaging, and quantitative analysis of optic nerve axons

Mouse optic nerves were collected, processed, stained with paraphenylenediamine (PPD), and imaged as previously described^32^. In brief, mouse heads were collected and submerged in half-strength Karnovsky fixative; nerves were dissected and embedded in resin, histologic sections were collected using an ultramicrotome (UC6, Leica, Wetzler, Germany) equipped with a diamond knife (Histo, Diatome, Hatfield, PA, USA), and stained with PPD, which stains the myelin sheath of normal axons and the axoplasm of dead or degenerating axons^33^. Stained optic nerve sections were imaged using a light microscope (BX52; Olympus, Tokyo, Japan) equipped with a camera (DP72, Olympus, Tokyo, Japan) and corresponding software (CellSens; Olympus, Tokyo, Japan). Before analyses, histological sections from each optic nerve were screened for quality control to ensure the plane of cutting was cross-sectional, sufficiently stained (i.e. areas within the tissue on the section that contained myelinated structures were PPD^+^, whereas areas outside the tissue were PPD^-^ and transparent), and the embedding resin was fully polymerized throughout the tissue. If these conditions were not met, sections were discarded and replaced.

Automated quantitative analyses were performed using Axon-Deep (*M. K. Garvin, personal communication, February 5, 2020*), an automated image analysis tool based on an algorithm that employs deep-learning to quantify optic nerve axons^34^. Manual axon quantifications were performed by an experienced technician masked to all other data, following previously described methodology^32^. Qualitative grading of optic nerve specimens was additionally performed by three skilled technicians masked to other data who used a three point optic nerve grading scale (increasing numerical grade with increasing damage: 1 = none to mild, 2 = moderate, 3 = severe) previously utilized to assess glaucomatous damage in mice^35^.

For validation of the Axon-Deep tool, automated counts by Axon-Deep from a subset of optic nerve specimens were compared to manual axon counts. Axons were quantified from a series of images (*n* = 80 images) also subjected to manual quantifications that were collected from optic nerves (*n* = 8 nerves, 1 nerve from each of 8 mice; 10 images per nerve) and compared using a Pearson’s Correlation Coefficient (r).

### Measurements of retinal area

Montages of whole retinas were generated by stitching together adjacent light microscopy fields (at a total magnification of 20X; using the same light microscopy setup referenced above) of H&E stained wholemounts using the Manual MIA function in CellSens image analysis software (CellSens; Olympus, Tokyo, Japan). All retinal montages were generated using identical microscope and camera settings. Area measurements were made by manually tracing the edges of stained retinal wholemounts using the polygon tracing tool and measure function contained in Fiji image analysis software^30^. Regions of retina containing artifacts resulting from the physical manipulation of the tissue, including folds, a common occurrence in the peripheral retina, and holes, were accounted for in the measurements of retinal area. Comparisons of retinal area by sex were done using an unpaired Student’s two-tailed *t*-test.

In addition to area measurements of retinas from J:DO mice, retinal area was also assessed from cohorts of inbred adult DBA/2J (4 mos. and 16 to 24 mos. of age) and C57BL/6J (2 to 5 mos. and 18 to 20 mos. of age) mice. Whenever possible, measurements were made from both retinas of each mouse studied. A subset of retinas from these cohorts were part of a previously published study^31^; in the current study, archived tissue underwent additional quantitative analyses of retinal area. Mean retinal area was calculated and compared between groups of different ages within the DBA/2J and C57BL/6J cohorts using an unpaired Students two-tailed *t*-test; comparisons between all three strains within the 2 to 5 months age group were made using an ANOVA with a Tukey’s post-hoc test for multiple corrections.

## Results

### Comparisons Relevant to GCC Thickness

In vivo retinal OCT images were collected (Fig. 1A) from a large cohort of J:DO mice and automated 3D segmentation was used to measure GCC thickness (Fig. 1B). Of 45 mice, all had overtly normal appearing retinas. Thickness of the GCC varied broadly between individuals (Fig. 1C; range 16.0 µm; mean ± SD, 62.4 ± 3.7 µm; coefficient of variation (CV) = 5.9). Within individuals, differences in GCC-thickness of the left and right eye were less varied (average ratio left/right = 0.99 ± 0.027; left: range 16.0 µm; mean ± SD, 62.0 ± 3.9 µm; CV = 6.3; right: range 15.5 µm; mean ± SD, 62.7 ± 3.5 µm; CV = 5.6) and were significantly correlated to one another (*r* = 0.89, P < 1.0E-4; Fig. 1C). Fundus exams showed many variations in pigmentation and vessel structure, but no indices of spotting characteristic of many retinal degenerations, nor overt abnormalities such as colobomas indicative of developmental anomalies (Supplemental Fig. 2). Following the in vivo analysis, retina and optic nerve tissues were collected for quantitative analyses of several cellular features (Fig. 2A-D). Ex vivo retinal measurements varied widely between individuals, with less variation between measurements made bilaterally in individual mice (Fig. 2E-F). Qualitatively, all histologic tissues appeared healthy, for example lacking fragmented nuclei in the retinal whole-mounts, and optic nerves lacking notable numbers of axons with dark axoplasmic staining by PPD. Using the damage grading scale for mice^35^, all optic nerves had a score of “1”—indicative of healthy nerves. As expected for mice with a diverse genetic background, the number of axons in the optic nerve varied widely (range = 24,339 to 69,517 axons; mean ± SD = 53,994 ± 6,495, *n* = 61 optic nerves from 31 mice; CV of 12.0%). These data indicate that J:DO mice are largely free of overt retinal disease through the ages tested and that there is a wide range of RGC-related phenotypic variation that is likely genetically determined.

**Figure 2.**
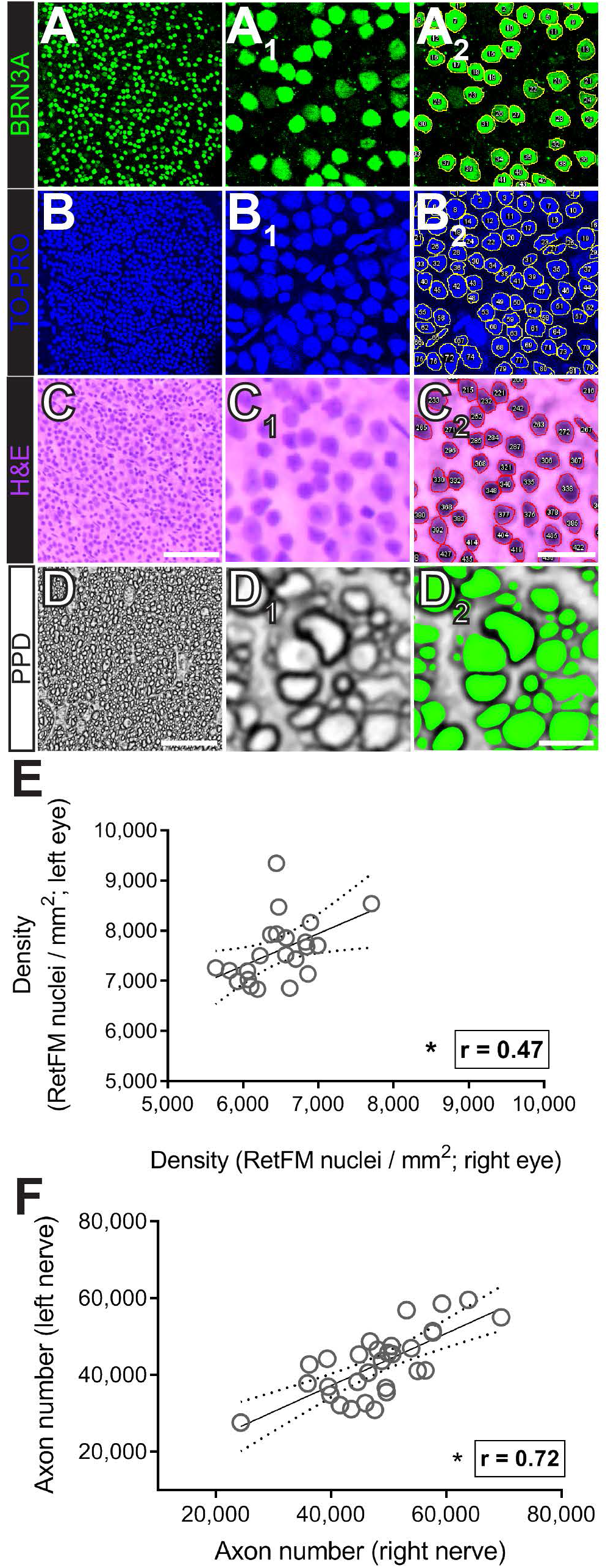
Quantitative analyses of neuronal features in retinas and optic nerves from J:DO mice. Representative micrographs of stained wholemount retinas and optic nerve cross-sections collected from adult J:DO mice used in quantitative analyses of neuronal features. Images in raw format (*first column, X*_*0*_) are of native size and magnification, whereas those in analyzed format are magnified and cropped down to better enable visualization of the neuronal features, both before (*second column, X*_*1*_) and after (*third column*; *features with inset contours or highlights, X*_*2*_) analyses. Confocal micrographs acquired from the same microscopy field of retina (**A**_**0-2**_) immunolabeled with an antibody targeting an RGC-specific marker (BRN3A, *in green*) and (**B**_**0-2**_) counterstained with TO-PRO (*nuclei, in blue*). Light micrographs from a (**C**_**0-2**_) retina stained with hematoxylin and eosin (*nuclei, violet; extracellular space, pink*) and (**D**_**0-2**_) optic nerve stained with para-phenylene diamine (PPD; myelin sheath, black). Scale bars = 100 µm (A_0_-C_0_), 25 µm (A_1,2_-C_1,2_), and = 10 µm (D_0_), and 2 µm (D_1,2_). Graphs relating quantifications of (**E**) total nuclei density (cells/mm^2^; *n*=22 mice) in the inner neural layers of retina and (**F**) axons (extrapolated number; *n*=31 mice) in optic nerve between the left and right nerves of the same individual amongst the population of J:DO mice. Each dot represents data from both retinas or nerves of one mouse, inset line represents best-fit line, Pearson’s correlation coefficient (r), and asterisks represent a P < 0.05 using a two-tailed Student’s *t*-Test.

In vivo and ex vivo measurements were then tested for correlations to one another (Table 1). Correlations between GCC thickness and average density of total nuclei in the inner retina were insignificant, whether measured in H&E-stained (Fig. 3A) or TO-PRO-labeled (Fig. 3B) samples. In contrast, correlations of GCC thickness with RGC-specific features were greater, including both average density of BRN3A^+^-nuclei (Fig. 3C) and number of PPD^+^ axons in the optic nerve (Fig. 3D); both correlations were statistically significant. Combined, these data indicate that among the variables we tested, GCC thickness correlates most closely with two metrics specific to RGCs, density of RGC soma in the retina and axon number in the optic nerve.

**Table 1.** Quantitative comparison of the relationships (Pearson’s Correlation Coefficients, r) between GCC-thickness and abundance of RGC somas in retina or RGC axons in optic nerve of the same eye and nerve pair from J:DO mice.

| Metrics correlated | $r =$ | $n =$ mice | $n =$ eyes or nerves | Significance | P- value |
| --- | --- | --- | --- | --- | --- |
| **GCC thickness with:** |  |  |  |  |  |
| Left versus right eye | 0.89 | 45 | 90 | * | < 1.0E-04 |
| BRN3A<sup>+</sup> nuclei density (left) | 0.52 | 28 | 28 | * | 7.7E-03 |
| TO-PRO<sup>+</sup> nuclei density (left) | 0.32 | 28 | 27 | n/s | 1.0E-01 |
| RetFM-J nuclei density (left and right) | 0.04 | 25 | 49 | n/s | 7.7E-01 |
| **Axon counts with:** |  |  |  |  |  |
| Left versus right optic nerve | 0.72 | 31 | 61 | * | < 1.0E-04 |
| GCC thickness (left and right) | 0.49 | 28 | 57 | * | < 1.0E-04 |
| - Left eye | 0.54 | 28 | 28 | * | 2.6E-03 |
| - Right eye | 0.51 | 28 | 29 | * | 5.0E-03 |
| NFL+GCL thickness (left and right) | 0.43 | 28 | 57 | * | 7.0E-04 |
| - Left eye | 0.36 | 28 | 28 | n/s | 5.7E-02 |
| - Right eye | 0.54 | 29 | 29 | * | 2.2E-03 |
| BRN3A<sup>+</sup> nuclei density (left) | 0.46 | 26 | 26 | * | 1.9E-02 |
| TO-PRO<sup>+</sup> nuclei density (left) | 0.45 | 24 | 24 | * | 2.7E-02 |
| RetFM-J nuclei density (left and right) | -0.03 | 25 | 46 | n/s | 8.3E-01 |

**Figure 3.**
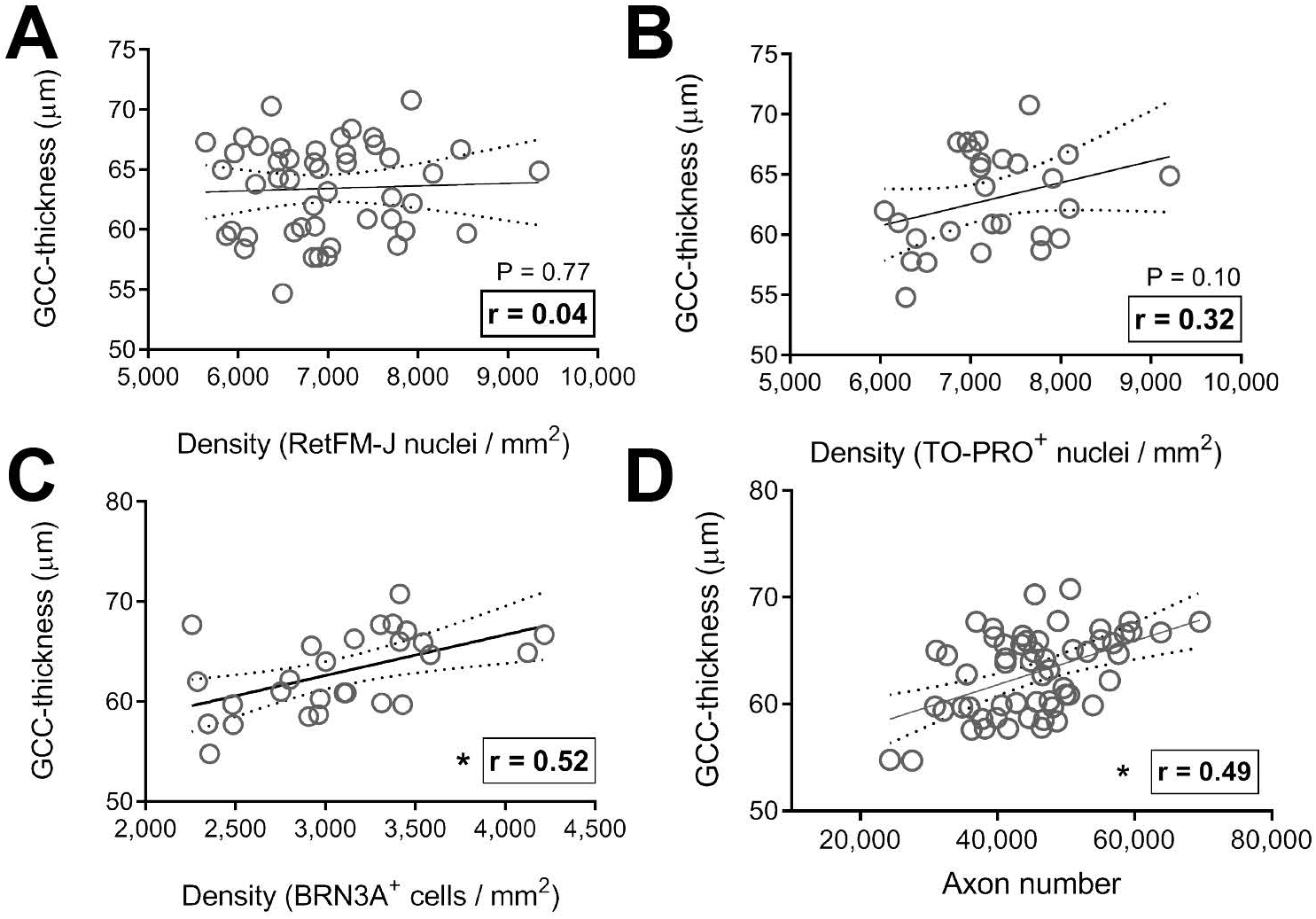
Relating cellular features to retinal structure in the ganglion cell complex of J:DO mice. Graphs relating how measurements of cellularity from retinal wholemounts relate to structural thickness by optical coherence tomography of ganglion cell complex (GCC) thickness in adult J:DO mice. Dot plots showing the relationship between structural thickness of the GCC (*y-axis*) versus the: overall cell density of all types residing in these layers, whether stained by (**A**) hematoxylin and eosin (*with analysis by RetFM-J*) or (**B**) TO-PRO (*with analyses using a custom macro for Image-J*), (**C**) density of retinal ganglion cells (*with BRN3A immunolabeling and analysis using a custom macro for Image-J*), or (**D**) extrapolated axon number. Note that the relationship between GCC thickness and overall cell density is relatively poor and not statistically significant, whereas those with RGC density and axon number are stronger and achieve significance. Each dot represents data from both eyes and/or nerves (*n*=47 for A and *n*=48 for D) or from one eye (*n*=27 for B and C) of one mouse, inset line represents best-fit line, Pearson’s correlation coefficient (r), and asterisks represent a P < 0.05 using a two-tailed Student’s *t*-Test.

A Fisher’s *Z*-Transformation was performed to compare the Pearson’s Correlation Coefficients for the strength of their relationships (Table 2). In these comparisons, GCC-thickness (*r* = 0.49) and BRN3A^+^-based methods (*r* = 0.46) for quantifying RGC soma correlated similarly to the axon count reference standard with no significant difference between the two techniques. A similar result was obtained when the same comparison (GCC-thickness: *r* = 0.33; BRN3A^+^ density: *r* = 0.45) was done relative to a smaller set of optic nerves (*n* = 8 nerves) that had also been manually counted.

**Table 2.**
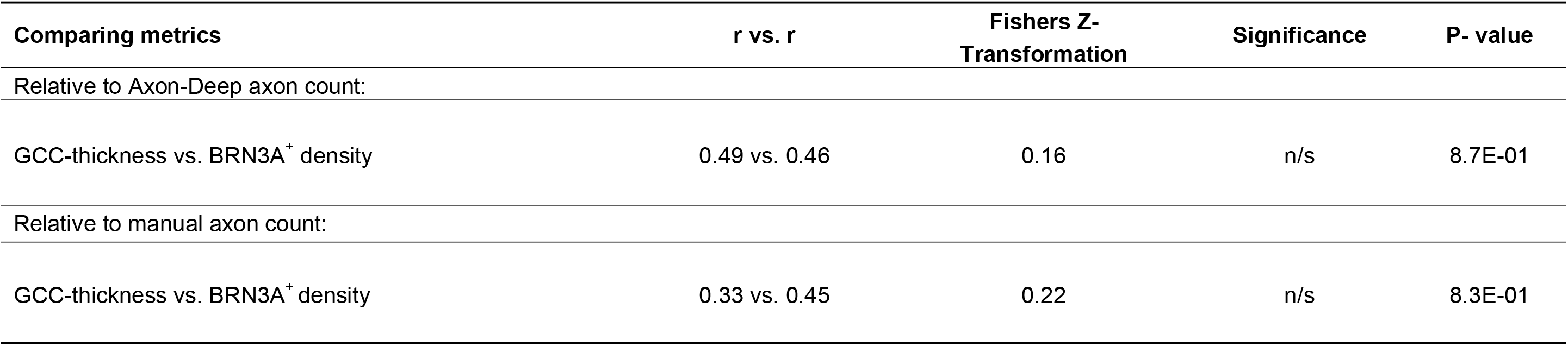
Comparing the relationship strength (Pearson’s Correlation Coefficient, r) between GCC-thickness and RGC somal counts to the manual axon count reference standard within the same eye and nerve pair from J:DO mice.

In our quantitative analyses of RGC metrics, the use of complementary assays in individual eyes provided opportunities for comparing the performance of some assays. Automated and manual axon counts were highly correlated (Supplemental Fig. 3, *r* = 0.94, P < 1.0E-4; *n* = 8 optic nerves, 1 from each of 8 mice), as were H&E- and TO-PRO-based counts of wholemount nuclei (Supplemental Fig. 4; central retina: *r* = 0.94, mid-peripheral retina: *r* = 0.87, peripheral retina: *r* = 0.67; P < 6.0E-4 for all three comparisons; *n* = 22 data pairs for each retinal zone from 22 retinas, 1 from each of 22 mice).

### Additional Comparisons Relevant to Retinal Anatomy and its Measurement

Variability in retinal size has previously been observed with buphthalmia secondary to elevated intraocular pressure^36^, amongst some strains with different genetic backgrounds^37^, and with natural aging^38, 39^—which we have replicated (Supplemental Fig. 5). In our analysis of the retinal wholemounts from these outbred J:DO mice, we observed that the size of the retina appeared to vary substantially (Fig. 4A-B). To quantify this, measurements from a subset of samples (*n* = 49 retinas, both from each of 27 mice) were made and total retinal area was indeed found to range from 15.0 to 26.1 mm^2^ (mean ± SD; 20.3 ± 2.4 mm^2^; Fig. 4C). Variation in retinal area was associated with changes in cellular density. Average densities of TO-PRO^+^ (Fig. 4D) and BRN3A^+^ nuclei (Fig. 4E) were both significantly correlated to retinal area, smaller densities being observed in retinas with larger area (*r* = −0.72, P < 1.0E^-4^; *r* = −0.65, *P* = 2.0E-4; respectively). Using total axon number as a surrogate for total RGC number, average density of BRN3A^+^ nuclei was also significantly correlated with RGC number (Fig. 4F), with density increasing as the number of axons increased (*r* = 0.46, *P* = 1.9E-2). Together, these observations point to a potential problem with common practices to quantify RGCs—two retinas with the same density of RGCs may or may not have a similar number of RGCs, depending on retinal area.

**Figure 4.**
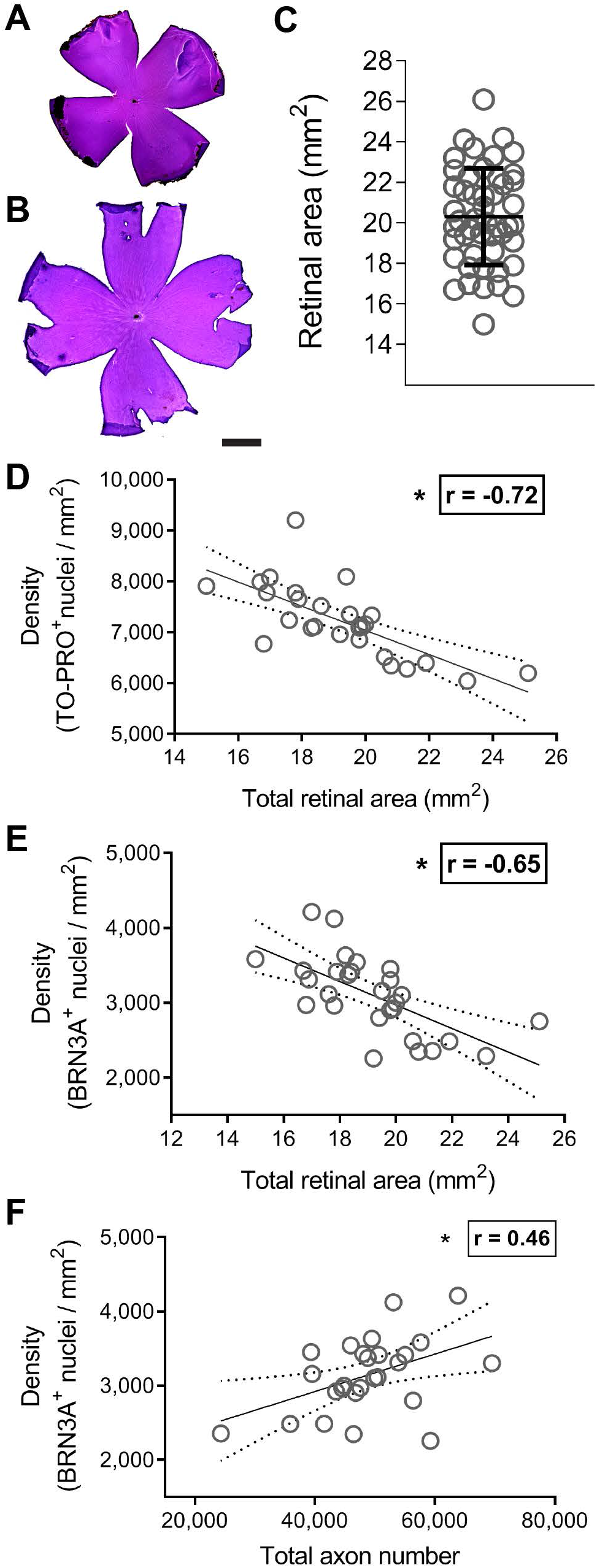
Relating cellular features in the ganglion cell complex to overall retinal area in J:DO mice. Image and graphical data relating how measurements of total area relate to cellular features collected from the same retinal wholemounts in adult J:DO mice. Representative light microscopy images of relatively (**A**) small and (**B**) large H&E-stained retinas collected from the J:DO cohort. Scale bar = 500 µm. (**C**) Graph showing the distribution of area measurements for all retinas (*n*=49) included in the study. Each dot represents data from one retina, inset horizontal and vertical lines represent the mean ± SD area for all retinas, respectively. Dot plots relating the: (**D**) density of TO-PRO^+^ nuclei and (**E**) density of BRN3A^+^ nuclei versus retinal area, respectively, and (**F**) density of BRN3A^+^ nuclei versus total axon number. Each dot represents data from the left eye/nerve pair from one mouse (*n*=27 pairs for D; *n*=28 pairs for E and F), inset solid and dotted lines represent the best-fit and 95% confidence interval, respectively, Pearson’s correlation coefficient (r), and inset asterisk to represent a P < 0.05 using a two-tailed Student’s *t*-Test.

As a complementary approach to ascertain total RGC number, we also calculated “estimated cellular number” purely from retinal data by multiplying the area of each zone of eccentricity in the retinal wholemounts by the average density in that same zone (Supplemental Fig. 6A-C), as done previously ^31^. Doing so resulted in a conversion of the average density of BRN3A^+^ nuclei from 3,117 ± 514 RGCS/mm^2^ to an extrapolated 53,946.5 ± 7,904.3 RGCs/retina. However, we did not detect a significant relationship between the estimated total number of BRN3A^+^ RGCs and total axon number (*r* = 0.20; *P* = 3.4E-1, Supplemental Fig. 6D), suggesting that this method of estimating total RGC number is, at present, relatively crude. This conversion to RGC number did not improve the strength of the correlations with GCC thickness (*r* = 0.42; *P* = 1.9E-2, Supplemental Fig. 6E).

Cross sectional area of the optic nerve also varied substantially between J:DO mice (ranging from 0.05 to 0.13 mm^2^; mean ± SD, 0.09 ± 0.02 mm^2^; *n* = 61 nerves; CV of 22%; Fig. 5A-G). There was a significant positive relationship between total cross-sectional area of the optic nerve and total axon number (*r* = 0.57; P < 1.0E-4; *n* = 61 optic nerves; Fig. 5H). Like the retina, increasing area of the optic nerve was associated with decreased average axon density (*r* = −0.43; P < 1.0E^-4^; *n* = 61 optic nerves; Fig. 5I). Average optic nerve cross sectional area and retinal area were not significantly associated with one another (Fig. 5J).

**Figure 5.**
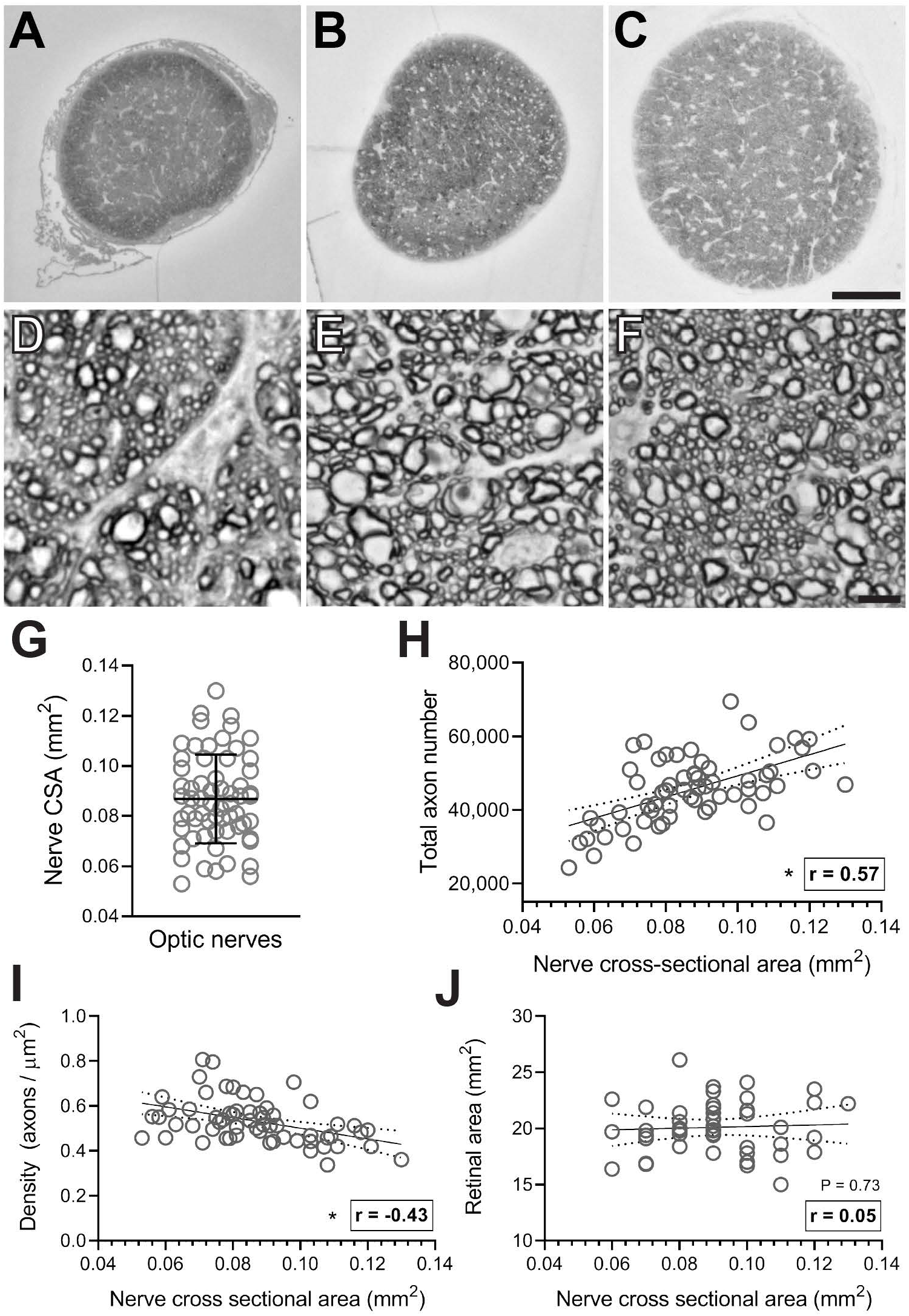
Relating variability in axon number with optic nerve structure in J:DO mice. Light micrograph pairs of para-phenylene diamine stained optic nerve cross sections of three representative optic nerves, presented in their entirety (*top row*) and magnified (*bottom row*), collected from adult J:DO mice. These micrographs were collected from optic nerves with the (**A, D**) smallest, (**B, E**) median, and (**C, F**) largest cross-sectional areas in the study cohort. Scale bars = 100 µm (100X total magnification; A-C) and 5 µm (1000X; D-F). (**G**) Graph showing the distribution of cross-sectional area (CSA) measurements for all nerves included in the study. Each dot represents data from one nerve, inset horizontal and vertical lines represent the mean ± SD area for all nerves, respectively. Dot plots relating (**H**) total axon number, (**I**) mean axon density, and (**J**) retinal surface area to the CSA of the corresponding optic nerve. Each dot represents data from one optic nerve (*n*=61) or nerve/retina pair (as in J; *n*=46), inset solid and dotted lines represent the best-fit and 95% confidence interval respectively, Pearson’s correlation coefficient (r), and asterisks represent a P < 0.05 using a two-tailed Student’s *t*-Test.

In the inbred mouse strains that have previously been studied, it is known that displaced amacrine cells constitute a significant fraction—approximately 50.3%—of the cells in the ganglion cell layer of mice^40^. However, it is unknown if this fraction is relatively constant or fluctuates according to genetic background. In our data, there was a significant positive relationship between the overall cell density of BRN3A^+^ and TO-PRO^+^ nuclei in J:DO retinas (*r* = 0.77, P < 1.0E-4, Supplemental Fig. 7A). BRN3A^+^ nuclei were 42.5 ± 4.7% (mean ± SD) of all TO-PRO^+^ nuclei, with a range from 32 to 52% (Supplemental Fig. 7B).

There were no significant differences in any parameters studied between male and female J:DO mice (Table 3).

**Table 3.**
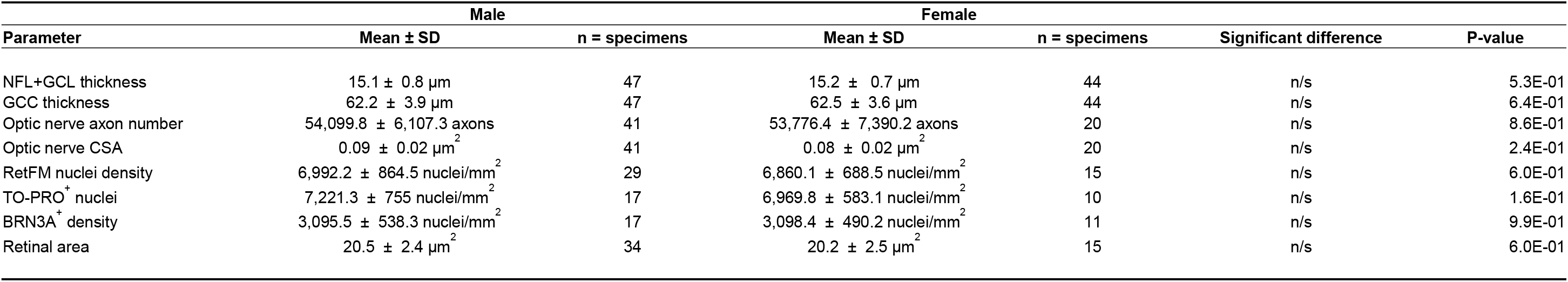
Cellular and structural features of retina and optic nerves in male versus female J:DO mice.

| Parameter | Male |  | Female |  | Significant difference | P-value |
| --- | --- | --- | --- | --- | --- | --- |
| | Mean ± SD | n = specimens | Mean ± SD | n = specimens | | |
| NFL+GCL thickness | 15.1 ± 0.8 μm | 47 | 15.2 ± 0.7 μm | 44 | n/s | 5.3E-01 |
| GCC thickness | 62.2 ± 3.9 μm | 47 | 62.5 ± 3.6 μm | 44 | n/s | 6.4E-01 |
| Optic nerve axon number | 54,099.8 ± 6,107.3 axons | 41 | 53,776.4 ± 7,390.2 axons | 20 | n/s | 8.6E-01 |
| Optic nerve CSA | 0.09 ± 0.02 μm<sup>2</sup> | 41 | 0.08 ± 0.02 μm<sup>2</sup> | 20 | n/s | 2.4E-01 |
| RetFM nuclei density | 6,992.2 ± 864.5 nuclei/mm<sup>2</sup> | 29 | 6,860.1 ± 688.5 nuclei/mm<sup>2</sup> | 15 | n/s | 6.0E-01 |
| TO-PRO<sup>+</sup> nuclei | 7,221.3 ± 755 nuclei/mm<sup>2</sup> | 17 | 6,969.8 ± 583.1 nuclei/mm<sup>2</sup> | 10 | n/s | 1.6E-01 |
| BRN3A<sup>+</sup> density | 3,095.5 ± 538.3 nuclei/mm<sup>2</sup> | 17 | 3,098.4 ± 490.2 nuclei/mm<sup>2</sup> | 11 | n/s | 9.9E-01 |
| Retinal area | 20.5 ± 2.4 μm<sup>2</sup> | 34 | 20.2 ± 2.5 μm<sup>2</sup> | 15 | n/s | 6.0E-01 |

## Discussion

Quantification of RGC abundance is an important aspect in clinical management of several diseases, and in research, it is often the cornerstone of studies using mice to study diseases of RGCs such as glaucoma. OCT-derived measurements have become popular metrics, despite some ambiguity in their biological properties. In humans and larger animals, OCT can be used to measure several features relevant to RGC health, including GCC thickness, retinal nerve fiber layer (RNFL) thickness, and optic nerve head morphology, among others. The comparatively small size of the mouse eye challenges reliable measurement of some of these tissues, leading us and others to use GCC thickness. Here, we have characterized retinal features of outbred J:DO mice and experimentally tested the degree to which OCT-derived measurements of GCC thickness may be correlated to total cellular density in the RGC layer, RGC density, and axon number in the optic nerve. The results indicate that there are indeed significant—but imperfect—correlations between GCC thickness and RGC abundance across a wide range of values in mice. Our analyses also uncovered or confirmed several features relevant to the basic biology of RGCs and their quantification.

This study compared in vivo OCT measurements of GCC thickness to direct ex vivo measurements made with dissected tissues of the same eyes—an experiment that would obviously be impractical in humans. The results found confirmatory evidence that GCC thickness correlates positively to RGC abundance. While several studies analyzed mouse retinas with both OCT and RGC-specific immunostaining^41–44^, relatively few have tested the correlations between these measurements^5, 45^. Nishioka and colleagues studied correlations between RGC density determined by immunostaining with RBPMS and OCT-derived GCC thickness in mice with experimental autoimmune encephalomyelitis (EAE)^5^. After 8 weeks of disease, a context in which RGC density had decreased by 23.3% and the remaining RGC somas had shrunk in cross sectional area by 27.4%, there was a significant correlation with GCC thickness (*r*i=i0.759, *P*lJ<i0.0001), which was slightly improved if baseline GCC thickness was considered and “GCC thinning” used instead. Likewise, Ho and colleagues compared histologically determined numbers of total cells in the ganglion cell layer from sampling of retinal cross sections with OCT-derived measurements of GCC thickness in mice with experimental anterior ischemic optic neuropathy. Both assays showed an initial retinal swelling, which was followed by atrophy. At 4 weeks following injury, a time point in which 49.3% of cells in the ganglion cell layer had been lost, there was substantial correlation between cell numbers and GCC thickness (*r*^*2*^ = 0.74). The degree of correlation detected in these studies was higher than the analogous correlation detected in our current study (*r* = 0.52, *P* = 5.0E-3), likely because of differences in the experimental context— especially using inbred mice with induced disease versus outbred mice with only natural variation. It is also notable that both studies detected acute *increases* in GCC thickness during initial stages of disease, which only later gave way to *decreases* in GCC. With respect to methodology, this illustrates how different stages of a disease might confound interpretations based solely on GCC thickness. For example, if a disease process was not temporally synchronous, as in these induced models, then the concurrent swelling and tissue loss in slightly different areas of the inner retina might lead to off-setting changes in GCC thickness.

In humans, correlations with OCT-derived metrics in the macula have been studied using models that estimate macular RGC number ^46^. In a study of 77 healthy, 154 suspect, and 159 glaucomatous eyes, Zhang et al. found that average thickness of the ganglion cell plus inner plexiform layer at the macula (mGCIPL thickness) was significantly correlated with the estimated number of macular RGCs (*r*^*2*^ = 0.67; P < 0.001). One reason for the less than perfect correlation in this study, that was considered by the authors and is also relevant to our current study, is that the ganglion cell layer also has displaced amacrine cells that contribute to GCC thickness in a manner not directly related to RGC number. In mice, the percentage of displaced amacrine cells in the ganglion cell layer is substantial, with different approaches reporting that RGCs are 36.1% to 67.5% of the neurons in the ganglion cell layer^47, 48^. A thorough study by Schlamp et al. used retrograde labeling to derive a percentage of 50.3%, while the estimate from using BRN3A as a marker of RGCs yielded an estimate of 44.8% in inbred C57BL/6J mice^40^. Our current study found that 42.5% of the total cells in the ganglion cell layer were BRN3A^+^ RGCs, which is very similar to the findings of Schlamp et al.—especially considering that Schlamp used nuclear morphology to exclude some cells from the denominator (RGCs / total neurons) whereas we included all cells (RGCs / total cells). A relevant new finding of our study was that even with a segregating genetic background, RGC and total cell densities were significantly correlated to one another in the ganglion cell layer, with only modest variation in their ratio (Supplemental Fig. 7). However, our study detected only modest correlations between total cell density and GCC thickness, presumably indicating that the predominant contribution to GCC thickness is specifically from RGCs—a finding that is supported by the fact that the GCC contains not only their somas similarly to other cells, but also includes the dendrites and axons of RGCs. Therefore, mouse studies performed in a context in which genetic backgrounds are not matched should expect that RGC abundance, total cell abundance (including displaced amacrine cells), and GCC thickness would all be likely to vary substantially at baseline in different mice, but that longitudinal studies of GCC thickness over time would predominantly reflect changes to RGCs.

The J:DO stock was generated by crosses between 144 early generation recombinant inbred lines contributing to the Collaborative Cross^49^, and thus incorporates similar genetic variation, including variation from all of the major phylogenetic branches present in laboratory mice (including wild-derived CAST, PWK, WSB, and five additional standard inbred strains, such as C57BL/6J). The genetic diversity of the stock can’t be maintained by traditional husbandry used in individual laboratories; the complex breeding scheme requires them to be acquired from The Jackson Laboratory. In general, J:DO mice were developed to promote genetic analysis of complex traits^50, 51^ and because it is sometimes desirable to perform studies in animals that, which like most humans, are not inbred^52, 53^. Our study shows that if overt retinal disease occurs in J:DO mice, it is likely rare (< 1 in 47 mice), but that there is broad range of variability in many retinal features which could be studied using quantitative genetic approaches. The finding that GCC thickness, total inner retinal cell density, and total axon number, were each highly correlated between the left and right tissues of individual mice, but varied widely between different J:DO mice, emphasizes that these traits likely have genetic underpinnings that could either be mapped, or exploited in gene expression studies as others have done in mouse studies of myopia^54^.

The finding that retinal and optic nerve cross sectional areas can naturally vary so dramatically has an important implication to studies that rely on measurements of RGC density, i.e. changes in retinal/optic nerve area and RGC number might be confused with one another. Because neither RGC number, nor cross sectional area of the retina/optic nerve are constant, a difference in density lacks meaning without a concurrent measurement of their tissue area. Because the variability of retinal cross-sectional area of outbred J:DO mice is large compared to inbred young DBA/2J or C57BL/6J mice (see Supplemental Fig. 5), it is likely that the variation in tissue sizes is genetic background dependent. Accordingly, any experiment in which the genetic background of cohorts being compared was not identical would be at risk for this confounding possibility. Some studies have used cross sectional area of the optic nerve as an indication of disease severity, which may be meaningful for models such as the DBA/2J model of glaucoma^55^, but our results caution against this approach for experiments in which genetic background is not isogenic. The apparent continuous nature of variability in tissue areas implies multigenic influences, making relevant genetic background matching between cohorts even harder. This same problem can also influence inbred mice, as illustrated by increases in retinal area that occur with natural aging (Supplemental Fig 5)^39^ and buphthalmia associated with elevated intraocular pressure in DBA/2 mice^36^—both of which alone could decrease RGC density and lead to exaggerated estimates of glaucomatous RGC loss. Based on these findings, we suggest that all studies of RGC abundance made using density measurements also report the tissue areas.

Aside from genetic background^56^, sex is another relevant biological variable that should be considered in mouse studies. Although sex could influence many other physiological or pathological events, we found that sex had no significant effects on the anatomical parameters of the inner retina measured in this study. Others have previously reported that there are sex-specific differences in retinal gene expression in mice, including in microglial and RGC-specific pathways^57^, and some glaucoma studies using mice have detected sex-specific differences in disease^58, 59^—which would seemingly be unrelated to the traits we have studied.

The current study has multiple caveats that warrant mention. First, although RGC abundance was measured in several complementary ways, the stains and antibodies used might have influenced some results. Axons were identified by the uptake of PPD, which stains the myelin sheath of healthy axons and the axoplasm of dead or degenerating axons. Due to its lipophilic nature, PPD only stains myelinated axons. Thus, our analyses were limited to myelinated axons and did not include unmyelinated axons. Likewise, the use of BRN3A as a marker of RGCs may have left some fraction of RGCs unlabeled. Among markers of RGCs, the reason we chose to use BRN3A is that it labels only the nucleus of RGCs, creating succinct regions of interest that automated imaging-based approaches can better distinguish in comparison to markers such as RBPMS, which is cytoplasmic^60^. The fraction of RGCs not labeled by BRN3A in rodents is not precisely known; studies based on immunostaining with rat wholemounts suggest it labels 96.2% of RGCs defined by retrograde labeling with FluoroGold (FG), if FG^+^ microglia are discounted^61^, or 87.9% if they are not^62^. In mice, various studies have reported that BRN3A labels 81.6% of RBPMS^+^ cells^60^ and 85.6% of FG^+^ cells^63^. More recent analyses utilizing single cell RNA sequencing have found that BRN3A-encoding RNA appears to be found in all sub-classes of mouse RGCs (albeit, at varying levels)^64, 65^. In sum, BRN3A appears to be present in a high percentage of RGCs, but any RGCs not detected by BRN3A immunolabeling could have confounded our analyses. Second, we have utilized semi-automated quantification approaches, which confer many advantages, but are undoubtedly still imperfect. While it is promising that the tools we are using can detect associations such as left-right eye correlations in individual mice, it’s possible that some associations have been blurred by a combination of methodologic shortcomings. Third, we anticipate that both OCT imaging for mice and approaches for automated quantifications of tissues will continue to evolve in the years to come. Thus, the confirmation that changes detected in RGCs via OCT are indeed significantly correlated with actual anatomical changes is important, but the degree of correlation currently detected may represent a “lower limit” which will improve through technological advances in the years to come. Finally, these analyses were performed in healthy mice with sources of variation that are natural, as opposed to disease-related. As illustrated by the discussion of EAE, it is likely that some disease processes will increase the discordance between OCT- and histologic-based measurements. The influence of age also remains to be tested.

In conclusion, we have characterized several quantitative phenotypes of RGCs, including GCC-thickness, density of RGCs in the retina, and axon number in the optic nerve, that exist amongst outbred J:DO mice. This initial characterization indicates that J:DO mice are free of overt retinal disease but have many ocular traits which vary widely between different individuals. Thus, J:DO mice are a powerful resource for studies such as ours—those that rely on maximal degrees of natural phenotypic diversity—and for future studies that employ quantitative genetic approaches to investigate these types of naturally occurring diversity that exist amongst individuals within a species. Of the phenotypic correlations tested, the most important finding was that non-invasive OCT-derived measurements of GCC thickness are significantly correlated with RGC abundance measured from histology-based analyses of retina and optic nerve. By extension, these results are consistent with a hypothesis that in human, OCT-derived measurements are likely also as valid as histologic quantifications of the retina. For mouse studies, our current results also indicate that tissue area is a potentially confounding variable in studies of RGC density; that the fraction of RGCs to displaced amacrine and other cells is relatively uniform, even if RGC abundance is not; and that there were no sex-specific differences in the RGC-related metrics measured.

## Acknowledgements

Work was supported by US Dept. of Veterans Affairs Rehabilitation Research and Development (RR&D, I01RX001481) and National Institutes of Health NEI (R01EY017673) grants to MGA. MDA is supported by the Robert C. Watzke endowed Professorship, and AHB was supported by Training Grant T32 DK112751-01. We also acknowledge an NIH/NEI Center Support Grant to the University of Iowa (P30EY025580). The contents do not represent the views of the U.S. Department of Veterans Affairs or the U.S. Government.

## Authors’ contributions

AHB collected all experimental data, performed histology and histological analyses, and spearheaded writing of the manuscript. KM conducted retinal OCT and fundus imaging. CJV performed the BRN3A/TO-PRO immunolabeling of retinas. KL measured retinal thickness from OCT images using the Iowa Reference Algorithms. DS measured retinal area. WD and MG quantified axons using the Axon-Deep tool. HM managed the mouse colony. MK and DP performed manual counts of optic nerve axons. KW assisted in the statistical analyses of data. All authors contributed to revision of the manuscript. MGA and MDA oversaw all aspects of this project.

