## Supplemental Figure Legends for "Biological Correlations and Confounding Variables for Quantification of Retinal Ganglion Cells Based on Optical Coherence Tomography using Diversity Outbred Mice"

**Supplemental Figure 1. Overview of the assays performed on retinal and optic nerve tissues from J:DO mice.** Vertical flow chart providing the numbers of mice (*n*=), tissue-types (retina or optic nerve) and -laterality (left or right) assessed, and the assays (in vivo imaging- or ex vivo histology) performed in the study of these tissues of J:DO mice at the indicated ages.

**Supplemental Figure 2. Variability in the appearance of the fundus and retinal vasculature across J:DO mice.** Montage of fluorescein angiogram images captured from a mixed cohort of albino and pigmented adult J:DO mice. Fundi of albino mice (*n*=6; **A**, **E**, **AF**, **AH**, **AK**, **AN**) exhibit a characteristic red-orange reflex, due to the absence of pigment in the eye, whereas fundi from pigmented mice exhibit a darker brownish reflex with prominent green fluorescent vessels (*n*=40; **B**-**D**, **F**-**AE**, **AG**, **AI**-**A**J, **AL**-**AM**, **AO**-**AT**). Phenotypic differences relate to variations in retinal vasculature, including vessel: number (for example: **AJ** versus **AL**), branching (for example: **V** versus **AS**), tortuosity (for example: **AQ** versus **AR**), and organization (for example: **J** versus **M**).

**Supplemental Figure 3. Validation of optic nerve axon quantification by Axon-Deep against ground truth manual quantifications.** Graph relating axon counts performed using Axon-Deep (*x-axis*) against a ground truth reference standard of axons counted manually (*y-axis*) from optic nerves of adult J:DO mice. Each dot represents data from one optic nerve from one mouse (*n*=8 nerves, 8 mice), inset solid and dotted lines represent the best-fit and 95% confidence interval respectively, Pearson’s correlation coefficient (r), and asterisk represents a P < 0.05 using a two-tailed Student’s *t*-Test.

**Supplemental Figure 4. Validation of the detection and quantification of nuclei in the inner retina by TO-PRO and subsequent H&E staining of the same retinas from J:DO mice.** Dot plots relating quantifications of total cellularity by zone of eccentricity, using two complementary methods (*TO-PRO and hematoxylin nuclear dyes followed by fluorescence and brightfield light microscopy, respectively*) applied to the same specimens in succession, in the inner retinal layers of wholemounts prepared from the left eyes of J:DO mice. The application of distinct nuclear dyes with independent detection methods in this approach provided a useful way to validate quantifications of cellularity in the (**A**) central, (**B**) mid-peripheral, and (**C**) peripheral zones of the inner retina. For all (A-C), each dot represents data from the left eye for one mouse (*n*=22), inset line represents best-fit line, Pearson’s correlation coefficient (r), inset asterisk to represent a P < 0.05 using a two-tailed Student’s *t*-Test.

**Supplemental Figure 5. Retinal area is variable in outbred J:DO mice, influenced by genetic background, and is dynamic with aging.** Graph showing measurements of total surface area of hematoxylin and eosin-stained retinal wholemounts collected from cohorts of adult outbred J:DO mice (5-mo-old; *n*=31 mice), inbred DBA/2J mice at ages before (4-mo-old; pre-glaucoma control; *n*=10 mice) and after (16 to 24 mos. of age; glaucoma control; *n*=9 mice) the onset of glaucoma, and inbred C57BL/6J mice (2 to 5 mos. and 18 to 20 mos. of age; normal control for aging; *n*=15 and 10 mice, respectively). Each dot represents data for one individual mouse, either from one retina or average between both (*when available*) retinas, inset lines represent mean area ± SD (horizontal and vertical, respectively), and inset asterisks represent a P < 0.05. Statistical comparisons to test the influence of age (*inset* *red lines and asterisk*) within inbred DBA/2J and C57BL/6J mouse strains were made using a Student’s t-test, whereas those testing the influence of genetic background (*inset* *black lines and asterisk*) amongst the three different mouse lines of similar age were made using a one-way ANOVA with Tukey’s post-hoc test for multiple corrections.

**Supplemental Figure 6. Variability in the structural and cell-based features along the retinal ganglion cell axonal complex in J:DO mice.** (**A**) Schematic of a retinal wholemount with subdivision of the three zones of eccentricity (*central, mid-peripheral, and peripheral zones delineated by dotted white lines; marked by increasingly dark shades of purple*) of retina to show how retinal area was measured and used for calculations of total retinal area, as well as total numbers of retinal ganglion and overall cell number from the inner retinal layers. The equation (*inset*) shows that extrapolations of total RGC number were calculated by summing the products of area and density (*A x ρ*) for each zone of eccentricity. Graphs showing quantifications of BRN3A^+^ nuclei expressed in terms of (**B**) density and (**C**) extrapolated total number. Each dot represents data from one eye (left side, *n*=27) for one mouse. Dot plots relating (**D**) total axon number (*n*=24 retinas) and (**E**) ganglion cell complex (GCC) thickness (*n*=25 retinas) to total BRN3A^+^ cell number within the same retinas. Each dot represents data from one eye and nerve pair (left side) for one mouse, inset line represents best-fit line, Pearson’s correlation coefficient (r), inset asterisk to represent a P < 0.05 or inset *p*-value if P > 0.05 using a two-tailed Student’s *t*-Test.

**Supplemental Figure 7. Relationships between retinal ganglion cells (RGCs) and non-RGC cell types in the inner retina of J:DO mice.** (**A**) Dot plot relating the density of BRN3A^+^ RGCs (*y-axis*) to overall density of TO-PRO^+^ nuclei (*x-axis*) in the inner retina. Inset solid and dotted lines represent the best-fit line and 95% confidence interval respectively, Pearson’s correlation coefficient (r), inset asterisk to represent a P < 0.05 using a two-tailed Student’s *t*-Test. (**B**) Horizontal graph showing the relative densities, expressed as a ratio of BRN3A^+^ RGCs to TO-PRO^+^ nuclei, in the inner retina of J:DO mice. Inset horizontal and vertical lines represent the mean ratio ± SD, respectively, for all retinas. For both (A-B), each dot represents data from one eye (*n*=27) from one mouse.
