## Supplementary figures and images for "Biological Correlations and Confounding Variables for Quantification of Retinal Ganglion Cells Based on Optical Coherence Tomography using Diversity Outbred Mice"

### Supplemental Figures

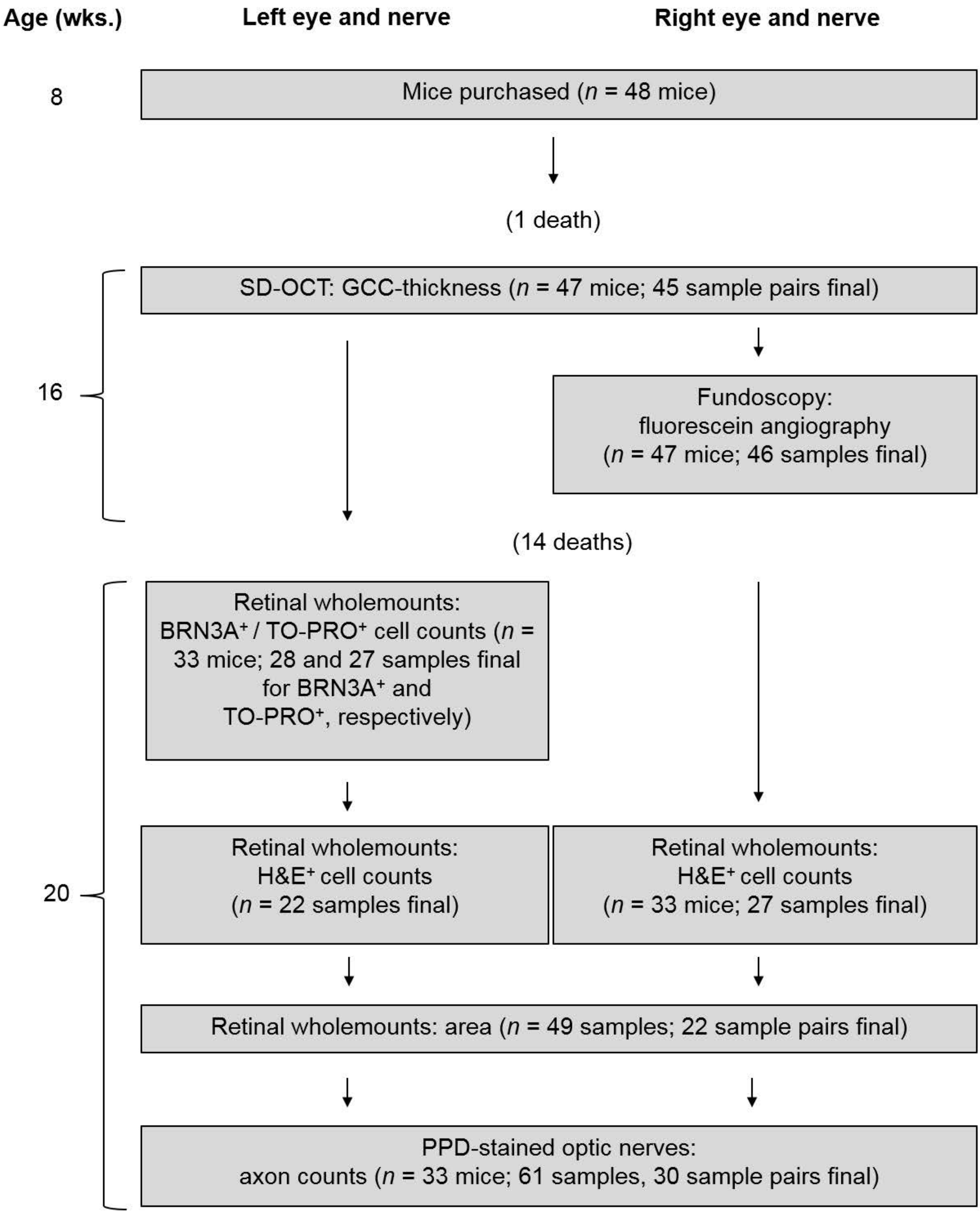

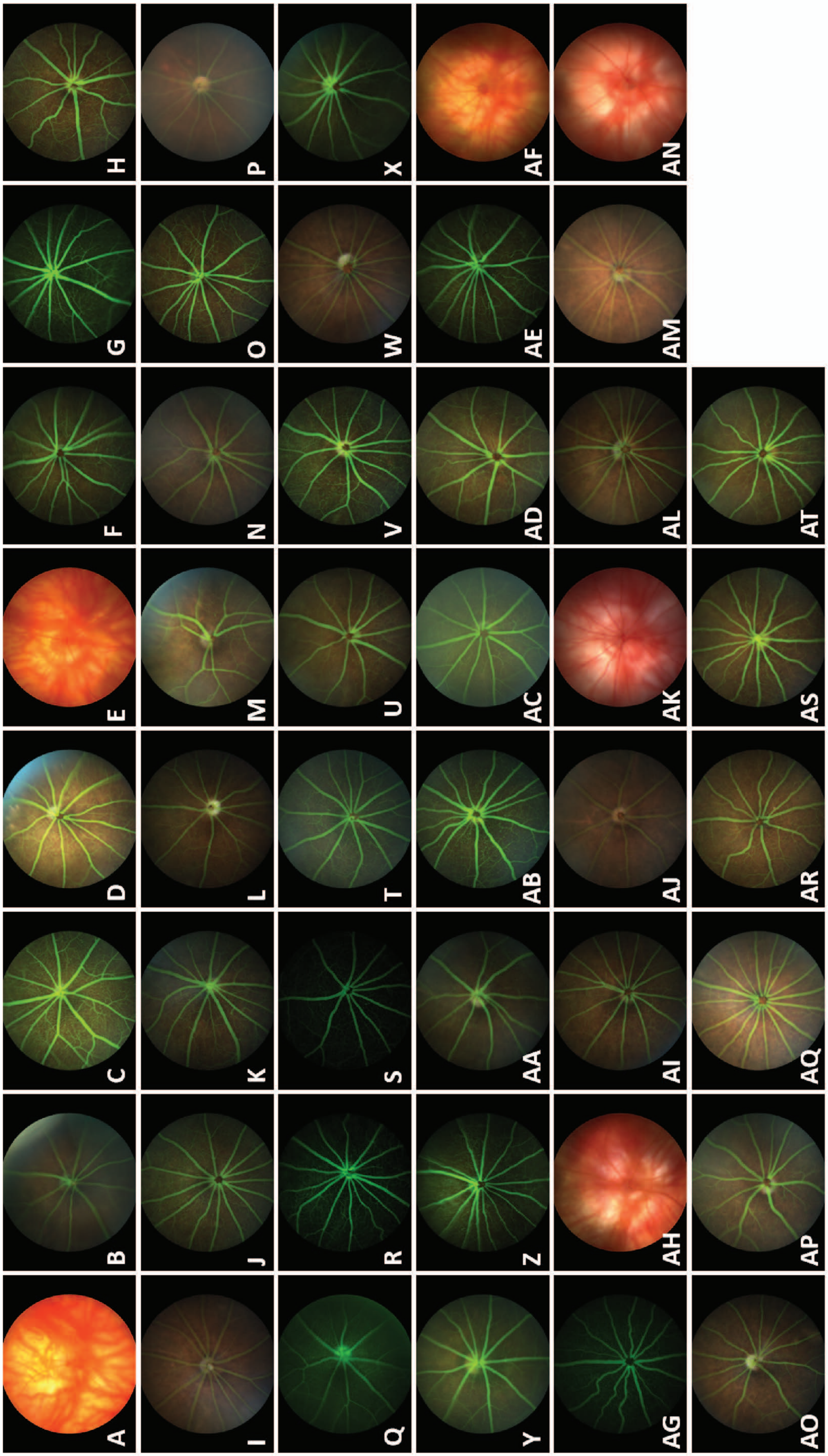

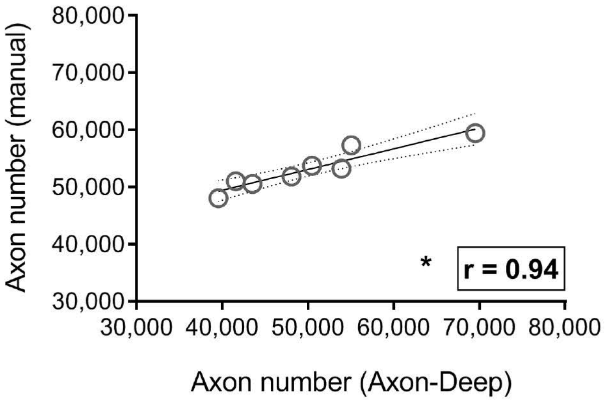

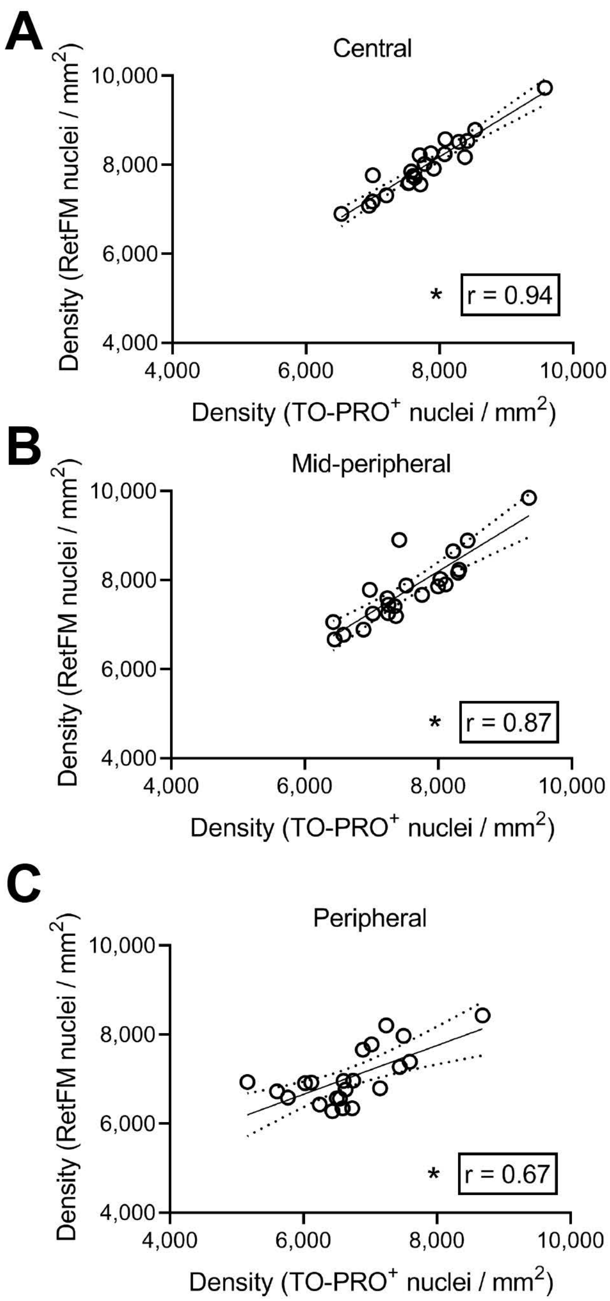

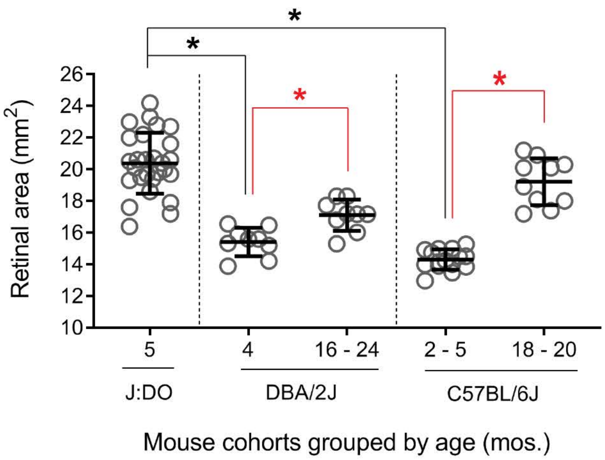

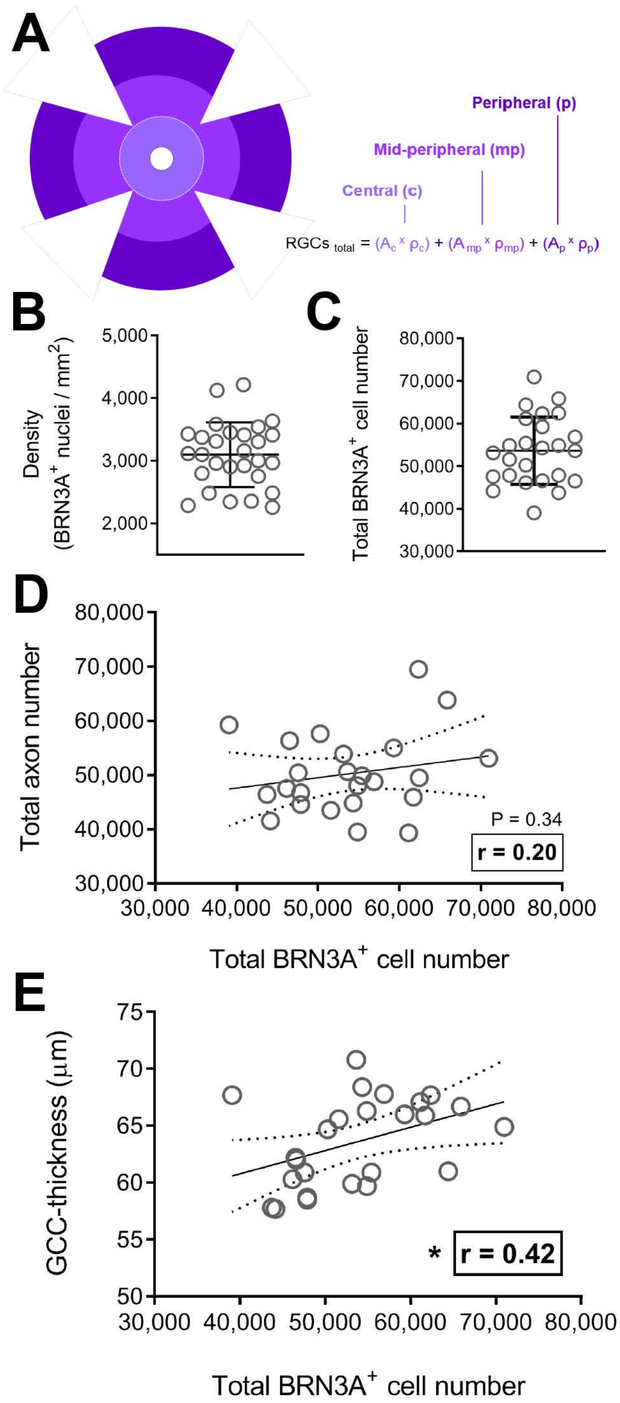

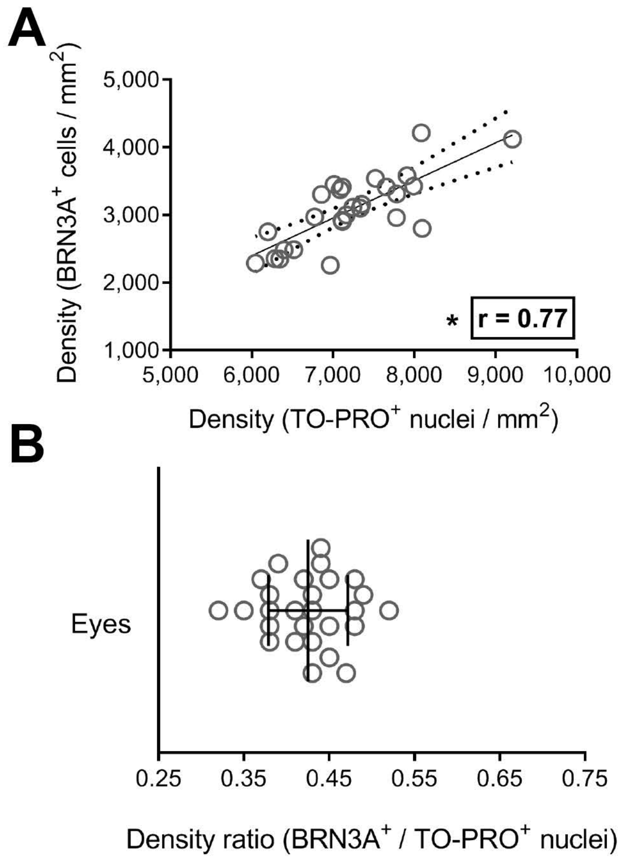
